# Physiological basis of photosynthetic hydrogen production in the cyanobacterium *Synechocystis*

**DOI:** 10.64898/2026.07.24.740500

**Authors:** Nadine Strabel, Florian Paul, Julia Regenbogen, Marko Boehm, Jens Appel, Kirstin Gutekunst

**Affiliations:** Molecular Plant Physiology, Bioenergetics in Photoautotrophs, University Kassel, Kassel, Germany

## Abstract

Photosynthetic hydrogen (photoH_2_) production by the cyanobacterium *Synechocystis* sp. PCC 6803 is an attractive means for storing solar energy. However, photoH_2_ yields remain limited by competing electron flux pathways. Recent *in vitro* characterization suggests that photoH_2_ production requires electrons from both carbohydrate oxidation and photosynthesis. Engineered fusions between photosystem I (PSI) and hydrogenase (PSI-H_2_ase) aim to divert electrons toward H_2_ production and rely exclusively on photosynthesis. Thus, photoH_2_ production differs fundamentally between wildtype (WT) and PSI-H_2_ase fusion mutants. Here, we show that photoH_2_ production in WT is enhanced by supplemented glucose, consistent with the recently reported confurcating nature of HoxEFUYH H_2_ases. PhotoH_2_ production was further studied in the new psaE-hoxUYH mutant by simultaneously monitoring electron flux through PSI alongside with turnover rates of O_2_, CO_2_ and H_2_. PsaE-hoxUYH achieved the highest photoH_2_ yield and longest production period among the currently available PSI-H_2_ase mutants in *Synechocystis*, prolonged by removing O_2_. Upon illumination, psaE-hoxUYH exhibited high initial photoH_2_ production rates, which decreased in parallel with CO_2_ fixation and ceased immediately in the presence of O_2_. In absence of O_2_, photoH_2_ production still declined slowly. Therefore, in addition to CO_2_ fixation and O_2_, other yet unknown factors might limit photoH_2_ production under these conditions. Moreover, we traced a previously observed high H_2_ production phase of unclear origin in psaD-hoxYH cultures to contaminating [FeFe]-H_2_ases from *Clostridium intestinale* rather than genuine photoH_2_ production by the mutant. Together, these findings indicate a complex metabolic interplay tuning photoH_2_ production in *Synechocystis* WT and PSI-H_2_ase fusion mutants.

## Introduction

The cyanobacterium *Synechocystis* sp. PCC 6803 (hereafter *Synechocystis*) possesses an oxygen sensitive bidirectional [NiFe]-hydrogenase (H_2_ase) HoxEFUYH. Under fermentative conditions, hydrogen (H_2_) is produced to balance the redox state of cells. Upon illumination of dark adapted, anaerobic cultures, a short burst of photosynthetic H_2_ (photoH_2_) is produced for a few minutes during which the CO_2_ fixation of the Calvin-Benson-Bassham (CBB) cycle is still inactive and O_2_ levels are low (1). Subsequently, photoH_2_ is consumed to recycle the electrons back into the metabolism (1, 2). In algae that possess [FeFe]-H_2_ases, photoH_2_ evolution was shown to cease due to competing electron demand by the CBB cycle before oxygen inactivates the enzyme (3-5). In cyanobacteria, NAD(P)(H) and ferredoxin_red/ox_ (Fdx_red/ox_) were discussed as redox partners for the HoxEFUYH H_2_ase for many years. PhotoH_2_ production was thought to be either based on Fdx_red_ or NADPH from photosystem I (PSI), linking photoH_2_ exclusively to electrons coming from the photosynthetic electron transfer chain (PETC) (1, 2, 6).

The recent purification of an intact and active HoxEFUYH holoenzyme allowed *in vitro* H_2_ production and consumption assays that revealed that NADP(H) does not function as redox partner for the enzyme. Instead, low H_2_ uptake rates were observed in the presence of just NAD^+^, while electron bifurcation utilizing NAD^+^ in combination with Fdx_ox_ resulted in high H_2_ uptake rates (7). H_2_ production strictly required both NADH and Fdx_red_ as electron donors in an elec-tron confurcating manner (7). These findings shed new light on the physiological role of photoH_2_ production in cyanobacteria *in vivo*. While Fdx_red_ is produced by PSI at the onset of light, NADH is formed during fermentative carbohydrate breakdown, linking photoH_2_ production to both processes (7). The idea of storing solar energy as photoH_2_ was particularly appealing, given that, in this process, electrons from photosynthesis are used to produce H_2_ without first passing through carbohydrate metabolism. Contrary to previous assumptions, the efficiency of converting solar energy into photoH_2_ in the WT appears lower than previously thought. However, this discovery supports the idea that using PSI-H_2_ase fusion mutants is a worthwhile approach for producing photoH_2_. In these mutants, the subunits HoxYH or HoxUYH of the native Hox hydrogenase are fused directly to PSI. According to current understanding, NADH transfers electrons via HoxF, and Fdx_red_ via HoxE to the H_2_ase (7). Since both subunits (HoxE and HoxF) are absent in the fusion mutants, electron transfer from NADH and Fdx_red_ should no longer be possible, so electrons for photoH_2_ production should come exclusively from PSI. One piece of evidence supporting this assumption is the observation that PSI-H_2_ase mutants are no longer able to take up H_2_, a process that would require an interaction with Fdx_ox_ and NAD^+^ (8-11). In *Synechocystis*, the H_2_ase (HoxYH or HoxUYH) has been attached to different PSI subunits (PsaC, PsaD, PsaE, PsaF) resulting in the mutants psaC-hoxYH, psaD-hoxYH, psaE-hoxYH, psaE-hoxUYH and psaF-hoxYH Δ*psaE*, including variants with different linkers between PSI and the H_2_ase (8-11). All listed PSI-H_2_ase mutants, except for psaC-hoxYH, produced photoH_2_. In psaF-hoxYH, the PSI subunit PsaE had to be deleted in addition, as it otherwise sterically hindered electron transfer between PSI and HoxYH (9). The first PSI-H_2_ase fusion mutant in *Synechocystis* was psaD-hoxYH (8). Two photoH_2_ production phases were reported for psaD-hoxYH and WT. The first transient phase was observed directly upon illumination reaching up to 3 μM H_2_ (WT) and 2.5 μM H_2_ (psaD-hoxYH). Under anoxic conditions in light, a second lasting H_2_ production phase was observed that commenced after several hours reaching concentrations between 30 μM H_2_ (WT) and 500 μM H_2_ (psaD-hoxYH) (8). However, tests on the light dependency of this higher H_2_ production had inconsistent effects. For instance, 3-(3,4-dichlorophenyl)-1,1-dimethylurea (DCMU), which blocks the electron transfer between PSII and the plastoquinone (PQ) pool, lowered H_2_ production in some cases as expected but enhanced the rates in others. Furthermore, it was unclear why the second H_2_ production period started with a delay of several hours, so it was initially assumed that a metabolic lag phase in *Synechocystis* might be responsible for this (8). In this study, we clarify the light dependency and the source of H_2_ production during the previously reported second H_2_ production phase in *Synechocystis* cultures. PsaD-hoxYH turned out to be unstable and to lose its photoH_2_ production capacity (8, 9). Attempts to generate new photoH_2_ producing psaD-hoxYH mutants have failed so far due to impaired strain growth. Therefore, the new psaE-hoxUYH mutant (11) was analyzed in detail. By simultaneous high-resolution measurements of H_2_, O_2_, and CO_2_ via membrane inlet mass spectrometry (MIMS) and parallel PSI electron flux measurements, the physiology and limiting factors of photoH_2_ production in WT and psaE-hoxUYH were studied further. Finally, our data show that photoH_2_ production by the native H_2_ase in *Synechocystis* WT depends on both photosynthesis and carbohydrate oxidation, which is well in line with *in vitro* data on the confurcating nature of HoxEFUYH (7).

## Results

### Light dependency of previously measured H_2_ concentrations in Synechocystis cultures

Hydrogen concentrations in long-term measurements had been monitored with H_2_ sensors from Unisense (Aarhus, Denmark) (8). The service lifetime of the sensors is limited, and their sensitivity decreases over time, which makes comparative measurements challenging. We therefore worked with a new set of sensors to collect reliable data. When we tested three of these new sensors in water under a polychromatic light regime alternating between 1 min darkness with 1 min light (light source: 12P HEX, ADJ, Thomann, Burgebrach, Germany), we observed that they all reacted with varying degrees of signal in response to light (Fig. 1B). This observed light sensitivity of the sensors impacts photoH_2_ measurements, as the burst of photoH_2_ production and uptake takes place within the first seconds to minutes after illumination. To avoid the light response, two approaches were tested in consultation with the company Unisense (Aarhus, Denmark). First, we tested different wavelengths (MULTI-COLOR-PAM, Walz, Effeltrich, Germany) and found that the sensitivity of the sensors to light was indeed dependent on wavelengths around 400 – 480 nm (Fig. 1A). Short-term photoH_2_ measurements that are routinely performed in monochromatic light at 625 nm in our lab (MULTI-COLOR-PAM and DUAL-KLAS-NIR, Walz, Effeltrich, Germany) are therefore not affected by the light sensitivity of the H_2_ sensors. In contrast, long-term photoH_2_ measurements, which were previously performed in experimental setups with polychromatic light, required improvements (8). Therefore, in a second approach, the sensors were painted black, which resolved the issue of their light sensitivity (Fig. 1C). The inconsistencies concerning the light dependence of H_2_ concentrations in previous long term measurements might therefore have been artifacts due to the varying light sensitivity of the utilized H_2_ sensors from Unisense (Aarhus, Denmark) (8).

**Figure 1:**
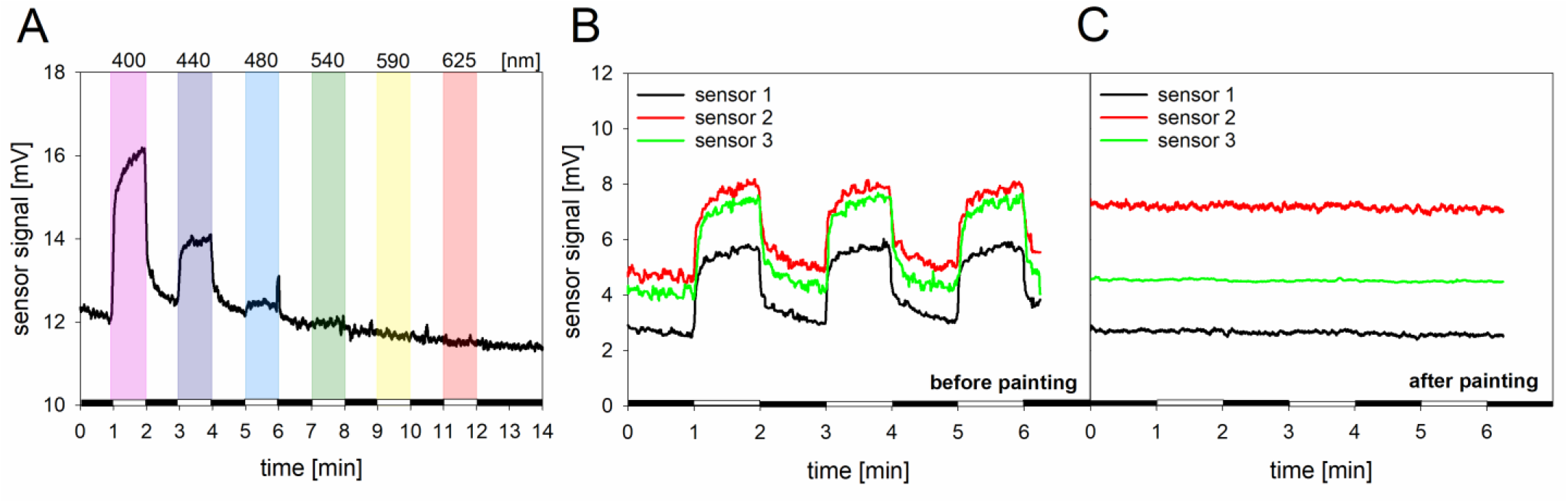
Light sensitivity tests of H_2_ sensors from Unisense (Aarhus, Denmark). **(A)** Test on the influence of different wavelength on the light sensitivity of sensors under monochromatic light as indicated. **(B-C)** Light sensitivity test in water under polychromatic light regime (light source: 12P HEX, ADJ, Thomann, Burgebrach, Germany) alternating between 1 min darkness and 1 min light, before and after painting sensor tip black as indicated.

### Contamination as source of previously reported high H_2_ yields in *Synechocystis* cultures

With the aim to clarify the physiology and source of lasting high H_2_ yields in *Synechocystis* cultures (8), respective experiments were repeated with WT accompanied by the D*hox* control strain, which contains a deletion of the entire hydrogenase *hoxEFUYH* operon. Anoxic conditions were achieved as before by the addition of glucose, glucose oxidase (GOX) and catalase (Cat) for O_2_ consumption. GOX/Cat reside in the culture medium, while as an undesired side effect, glucose can be taken up and metabolized by the cyanobacterial cells. Gluconate, a byproduct of this oxygen scavenging reaction, is not metabolized by *Synechocystis* and therefore resides in the culture medium (17).

When measuring long-term photoH_2_ with four WT and two Δ*hox* cultures without added antibiotics, three WT and both Δ*hox* cultures showed lasting H_2_ production reaching between 500-1500 μM H_2_ after a time delay of several hours (Fig. 2A). The experiment was repeated with six Δ*hox* cultures, three of which were treated with the antibiotic kanamycin. In this case, H_2_ production reaching between 800-1000 μM H_2_ was only observed in samples that did not contain the antibiotic (Fig. 2B). To check the cultures for potential contamination, three samples were taken from cultures (Fig. S2). For metagenomic analysis, one reference sample was taken before starting the experiment and two samples were taken during the H_2_ production phase. The reference sample contained 99.998% *Synechocystis*, while other organisms could be identified with abundances below 0.001% (Fig. 2C). The other two samples that were taken during the H_2_ production phase contained 99.867% and 99.807% *Synechocystis*, followed by 0.130% and 0.190% *Clostridium intestinale* respectively as second most abundant organism (Fig. 2D-E). Other bacteria were detected at significantly lower concentrations below 0.001%. Notably, *C. intestinale* could not be detected in the reference sample. The genome of *C. intestinale* possesses five [FeFe]-hydrogenases (18), three of which could be confirmed in both samples via PCR (Fig. 2F).

**Figure 2:**
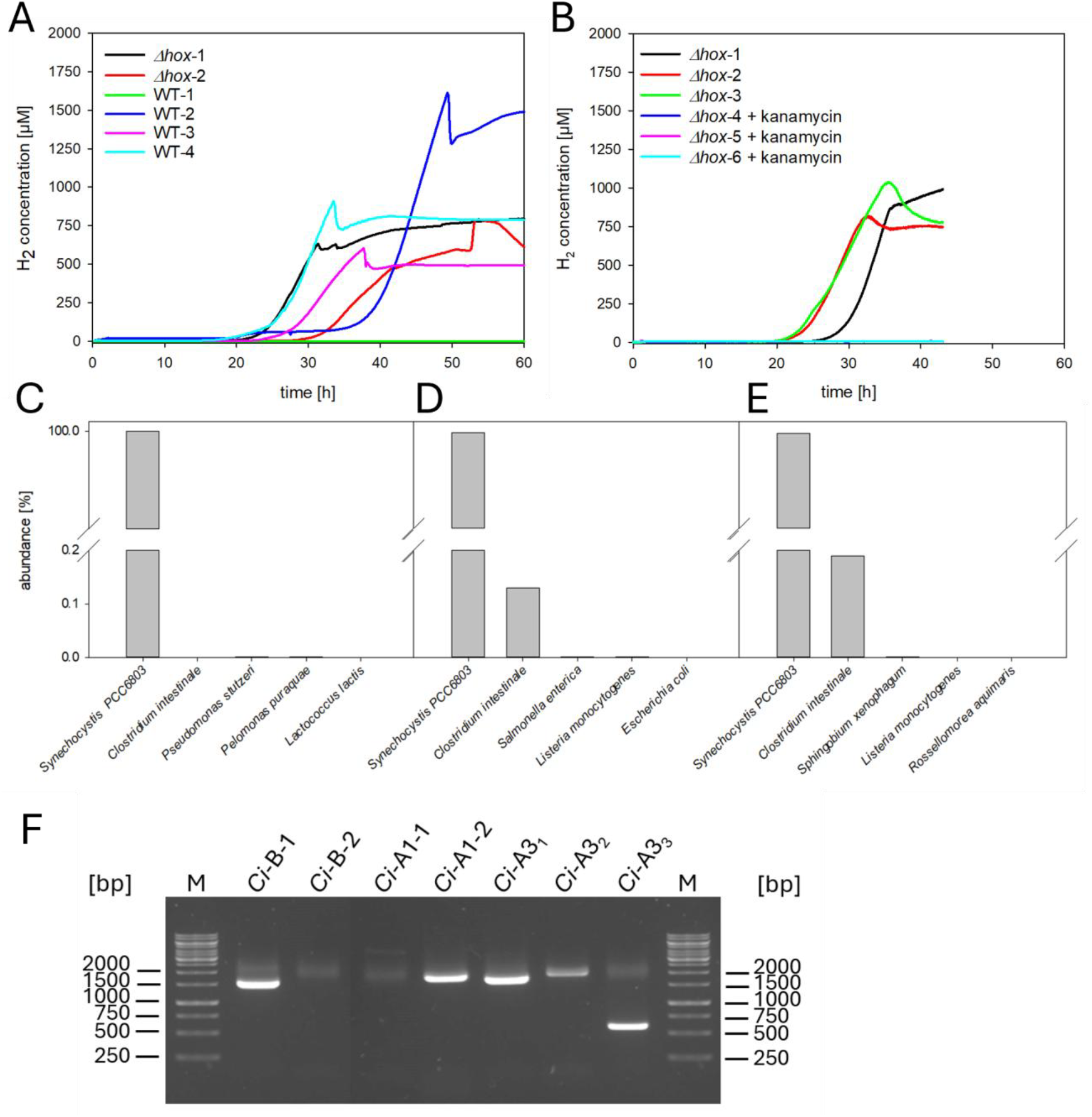
Source of high yield H_2_ production in Synechocystis WT and Δ*hox* cultures under anoxic conditions (10 mM glucose, 40 U/ml GOX, 50 U/ml Cat) in light. **(A)** H_2_ concentration in four WT and two Δ*hox* cultures without antibiotics. **(B)** H_2_ concentration in six Δ*hox* cultures, three were supplied with 50 μg/ml kanamycin. **(C-E)** Organismic composition of the cultures before (C) and during H_2_ production **(D, E)** as indicated in Fig. S2, according to metagenome analyses. F: PCR confirming the genetic presence of three [FeFe]-hydrogenases from *Clostridium intestinale* (Ci). Out of the genes from five hydrogenases investigated, Ci-B-1, Ci-B-2, Ci-A1-1, Ci-A1-2 and Ci-A3 (heterotrimer), three could be verified. Samples Ci-B-2 and Ci-A1-1 yielded no significant PCR products. The abbreviations B, A1 and A3 refer to the classification of the respective [FeFe]-hydrogenases.

The light sensitivity towards polychromatic light of H_2_ sensors from Unisense (Aarhus, Denmark) further complicates the interpretation of previous data. However, as photoH_2_ production in psaD-hoxYH was also measured in monochromatic light yielding up to 3.5 μM H_2_, the lower photoH_2_ levels that were reached in current measurements reaching around 0.2 μM H_2_ can be assumed to be caused by an instability of the psaD-hoxYH mutant with regard to photoH_2_ production (9).

### Contribution of glucose supplementation to fermentative and photoH_2_ production in the WT

To prevent inhibition of the H_2_ase, glucose and GOX/Cat were added to experiments for O_2_ consumption. As glucose is metabolized by *Synechocystis*, an alternative O_2_ scavenging system was tested based on ethanol, alcohol oxidase (AOX) and Cat. AOX oxidizes ethanol and O_2_ to acetaldehyde and H_2_O_2_. Cat subsequently splits 2 H_2_O_2_ to 2 H_2_O and O_2_ (19). Fermentative and photoH_2_ production were significantly impaired in the presence of ethanol and AOX/Cat, which might be due to the toxicity of the formed acetaldehyde (Fig. 3C-D). Therefore, the O_2_ scavenging system consisting of glucose and GOX/Cat is preferable. In order to estimate the contribution of externally added glucose to fermentative and photoH_2_ production, a hexokinase (HK; sll0539) deletion mutant (D*hk*), which is blocked in its ability to metabolize supplemented glucose, was characterized (17, 20). As previously mentioned, H_2_ production with HoxEFUYH requires both NADH and Fdx_red_ *in vitro* (7). *In vivo*, the main sources for Fdx_red_ are the pyruvate:ferredoxin oxidoreductase (PFOR) and PSI, while the main source for NADH is most likely glyceraldehyde-3-phosphate dehydrogenase GAPDH1 (Slr0884). GAPDH1 and PFOR are both involved in glucose oxidation. Glucose originates either from supplemented glucose or from the breakdown of internal glycogen reservoirs.

**Figure 3:**
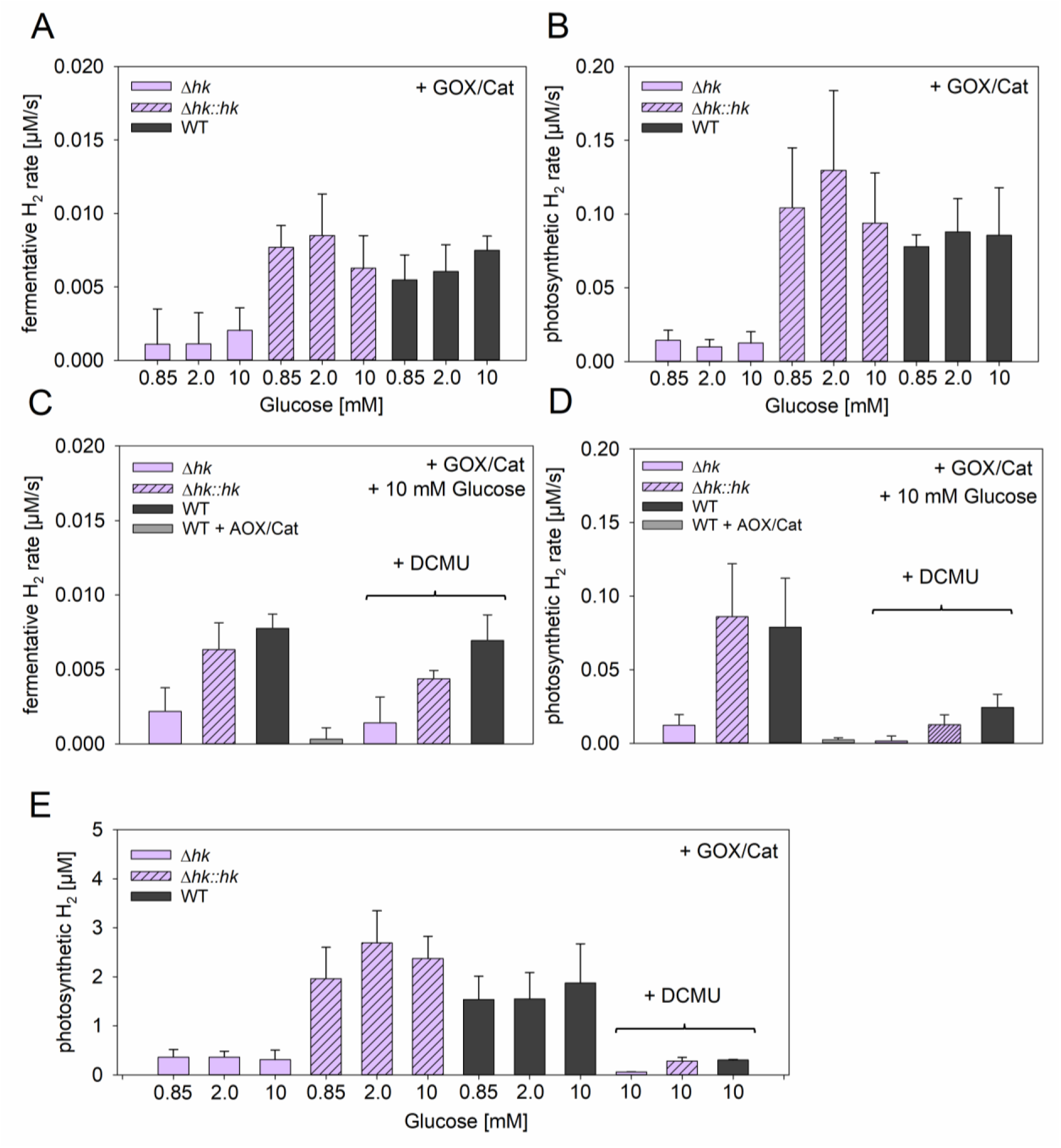
Fermentative and photosynthetic H_2_ production rates and concentration in Δ*hk*, Δ*hk::hk* and WT. Cells were adapted to darkness for 10 minutes, followed by saturating (∼800 μmol photons m^-2^ s^-1^, 625 nm) illumination for 10-20 minutes. The displayed rates are maximum H_2_ production rates during fermentative (dark) and photosynthetic (light) conditions. Enzyme mix (GOX/Cat: 40 U/ml GOX, 50 U/ml Cat) and 0.85-10 mM glucose were added for O_2_ scavenging. **(C-E)** 40 μM DCMU was added to inhibit electron transfer between PSII and plastoquinone (PQ). An alternative oxygen-scavenging system of AOX/Cat (AOX/Cat: 20 U/ml AOX, 25 U/ml Cat) and 2 mM ethanol was tested as well.

In Δ*hk*, which cannot utilize supplemented glucose, fermentative H_2_ production rates decreased to 32% and photoH_2_ production rates decreased to 15% compared to the WT (Fig. 3A-B). The photoH_2_ yield was reduced to 13% (Fig. 3E). The observed phenotypes of Δ*hk* could be complemented in Δ*hk::hk* (Fig. 3A-E). These data show that external glucose strongly contributes to both fermentative and photoH_2_ production rates and yield, while glycogen degradation supports photoH_2_ production at low rates and yield.

We next tried to estimate the contribution of H_2_O splitting at PSII to photoH_2_ production by adding DCMU, which blocks electron transfer between PSII and the plastoquinone (PQ) pool. In this setup, electrons from glucose oxidation can still be fed into the photosynthetic electron transfer chain via the NDH-1 complex and end up at PSI for Fdx_red_ production. Fermentative H_2_ production was not significantly influenced by DCMU, as expected. However, the residual photoH_2_ production rate and photoH_2_ yield in Δ*hk* were reduced close to zero by DCMU. In WT and Δ*hk::hk*, DCMU reduced photoH_2_ production rates to 37% and 28% and photoH_2_ yields to 16% and 12% respectively. Our data does not allow to quantify the exact contribution of electrons originating from either H_2_O or glucose oxidation to photoH_2_ production. It is difficult to discriminate between internal and external glucose oxidation, and addition of DCMU potentially enhances feeding of electrons from glucose oxidation into the photosynthetic electron chain, thereby distorting the original situation. However, our results clearly show that photoH_2_ production *in vivo* can be supported by glucose oxidation in absence of PSII activity. PhotoH_2_ production in the WT is most likely supported by a combination of photosynthesis and carbohydrate oxidation, which is in line with *in vitro* data on the confurcating nature of HoxEFUYH (7). These results emphasize PSI-H_2_ase fusion mutants as promising candidates for photoH_2_ production that should be exclusively linked to photosynthesis.

### Construction and characterization of the PSI-H_2_ase fusion mutant psaE-HoxUYH

With the aim to further improve photoH_2_ yields in PSI-H_2_ase fusion mutants and to investigate the physiology of photoH_2_ production in detail, psaE-HoxUYH was generated (11). PSI subunit PsaE was chosen as an anchor for the hydrogenase, as the previously constructed psaE-HoxYH mutant grew better than psaD-HoxYH and turned out to be stable with regard to photoH_2_ production (10). The electron transfer rates between PSI and HoxYH in previously constructed PSI-H_2_ase fusion mutants in *Synechocystis* are low, which might be attributed to non-ideal distances of more than 14 _Å_ between the FeS cluster F_B_ in PSI and the FeS cluster in HoxY (8, 10). Therefore, to improve electron transfer, the diaphorase subunit HoxU was integrated in psaE-HoxUYH since this subunit contains several FeS clusters (U1-U4) (11). The construct of psaE-HoxUYH was introduced into a Δ*hox*Δ*psaE* background strain, resulting in the mutant Δ*hox*Δ*psaE*::*psaE-hoxUYH*, which will be referred to as psaE-HoxUYH. The respective fusion strategy is shown in Fig. 4A-B. In parallel to this study, this mutant was also tested for photoH_2_ production when embedded in a viologen-modified redox polymer for O_2_ consumption without detailed mutant characterization (11).

**Figure 4:**
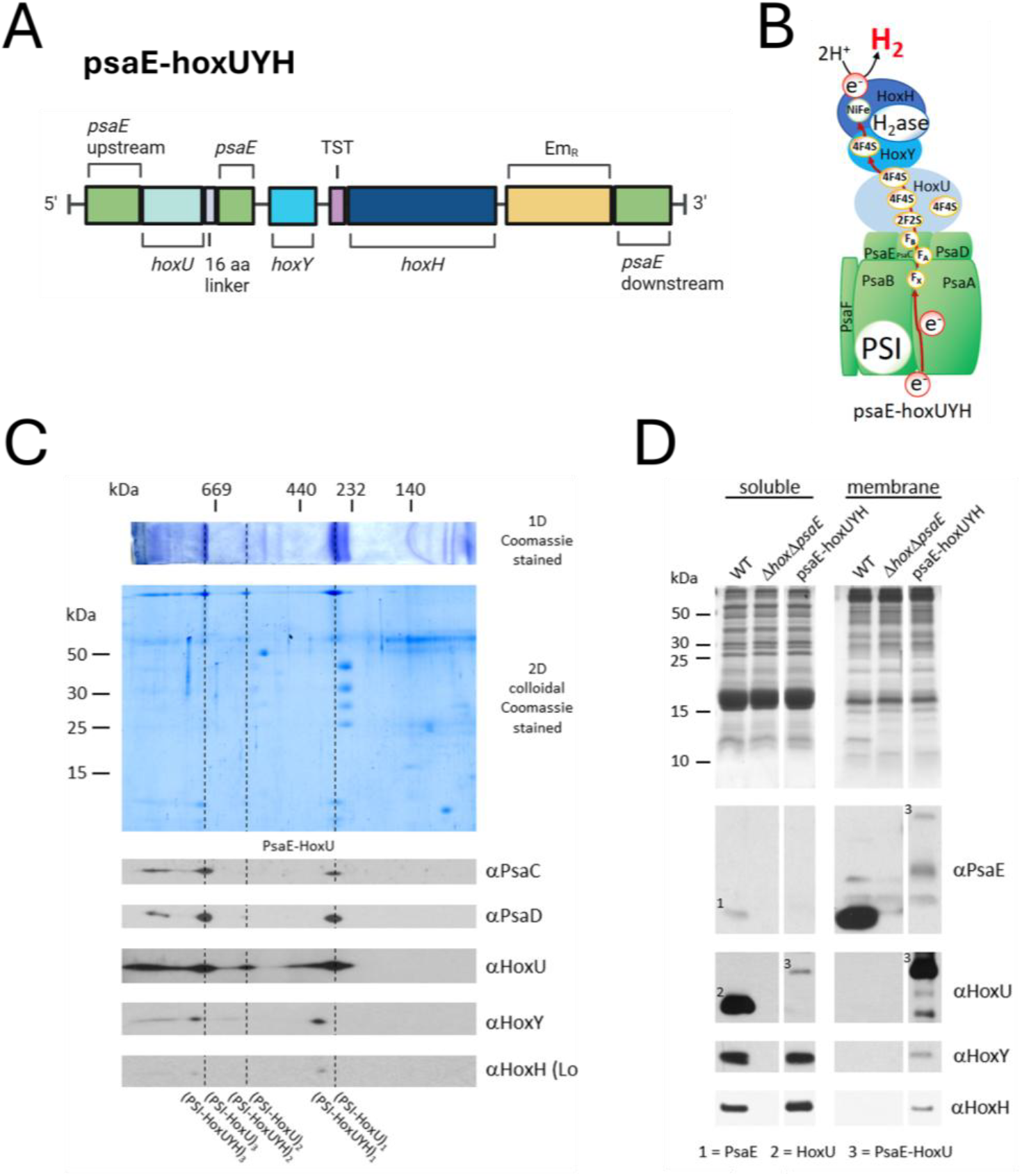
Fusion strategy, schematic structure and immunoblot of psaE-hoxUYH complexes. **(A)** The fusion construct is designed for homologous recombination into the psaE locus in a Δ*hox*Δ*psaE* background strain. PsaE is covalently fused to HoxU via a 16 amino acid linker and HoxH is N-terminally fused with a twin-strep-tag. **(B)** Schematic structure of psaE-hoxUYH fusion complex, **(C)** One-dimensional (1D) Blue Native PAGE, two-dimensional (2D) SDS-PAGE and Immunoblot of psaE-hoxUYH. Protein gels were stained with Coomassie. 1D-PAGE yielded different complexes that were separated into a second dimension with SDS-PAGE, revealing their subunit composition. Via immunoblotting with specific antibodies, the presence of PSI subunits PsaC and PsaD as well as Hox subunits HoxU, HoxY and HoxH were investigated. Three identified PSI complexes (monomeric, dimeric and trimeric) are indicated with a dotted line. **(D)** Coomassie-stained 1D-SDS-PAGE and Immunoblot of psaE-hoxUYH in comparison to WT, Δ*hox*Δ*psaE* and psaE-hoxYH in soluble and membrane fractions.

Immunoblots (Fig. 4C) show three complexes containing PSI as well as HoxU (PSI-HoxU)_1-3_, while Hox subunit signals HoxY and HoxH are shifted to a higher complex size (PSI-HoxUYH)_1-3_. Thus, the expression of the complete PSI-H_2_ase complex could be confirmed. In psaE-hoxUYH, signals for PsaE and HoxU were shifted to a higher molecular size compared the WT, confirming that both proteins are covalently linked (Fig. 4D).

Subsequently, psaE-hoxUYH was compared with WT and psaE-hoxYH (10) concerning photoautotrophic growth, H_2_ase activity, fermentative and photoH_2_ production (Fig. 5). Photoautotrophic growth of psaE-hoxUYH appeared similar to psaE-hoxYH and the WT (Fig. 5A). The expression levels of functional H_2_ase in both fusion mutants were about 30-40% lower than in the WT (Fig. 5B). Under fermentative conditions, only the WT strain produced H_2_ since both fusion mutants lack the HoxEF subunits which are required for the interaction with the natural redox partners Fdx_red_ and NADH (Fig. 5C, 5E) (7). Upon illumination, WT showed fast photoH_2_ production followed by H_2_ uptake. Both mutants did not take up H_2_, and their production rate was notably lower compared to the WT. However, the implementation of the HoxU subunit in psaE-HoxUYH increased the maximum photoH_2_ production rate 3-fold compared to psaE-HoxYH (Fig. 5E). Also, in comparison to WT and psaE-hoxYH, the duration of photoH_2_ production and yield of photoH_2_ were highest in psaE-hoxUYH (Fig. 5F, 5G). Therefore, psaE-hoxUYH appeared as a promising candidate for further characterization.

**Figure 5:**
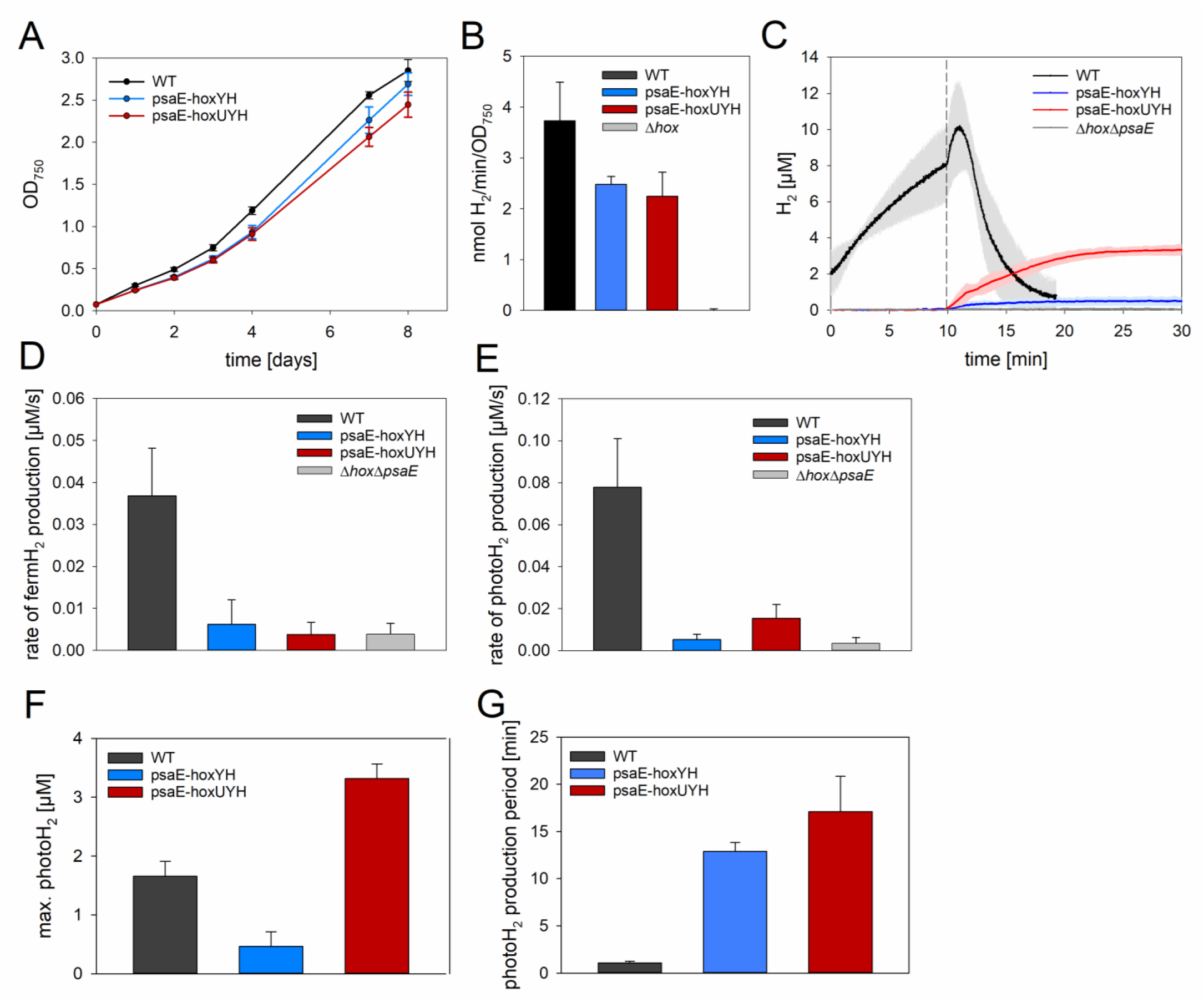
Photoautotrophic growth, hydrogenase activity and photoH_2_ production of *Synechocystis* WT, psaE-hoxYH and psaE-hoxUYH. **(A)** Photoautotrophic growth in BG-11 media at 50 μmol photons m^-2^ s^-1^ for 8 days (n = 3). **(B)** Methylviologen assay to determine hydrogenase activity with 10 mM sodium dithionite and 5 mM methyl viologen using a Clark-type electrode (S1 disc, Hansatech Instruments Ltd) (n = 3). **(C)** Fermentative (fermH_2_) and photoH_2_ production using a Clark-type H_2_-sensor from Unisense under anoxic conditions (10 mM glucose, 40 U/ml GOX, 50 U/ml Cat) (n = 3); cells were adjusted to a chlorophyll concentration of 20 μg/ml. After 10 minutes of darkness, cells were exposed to saturating light (∼800 μmol photons m^-2^ s^-1^, 625 nm) to induce photoH_2_ production. **(D-E)** Respective photosynthetic and fermentative H_2_ production rates (n = 3). **(F)** Maximum photoH_2_ production yield (n = 3). (G) Duration of photoH_2_ production (n = 3).

### Physiology of photoH_2_ production in Synechocystis WT and psaE-HoxUYH

To better understand the physiology and limiting factors of photoH_2_ production in WT and psaE-hoxUYH, concentrations of O_2_, CO_2_, H_2_ and electron flux through PSI were monitored in parallel (Fig. 6). Anoxic conditions were achieved with GOX/Cat and different concentrations of glucose to vary the duration of oxygen depletion. In dark conditions, all strains showed CO_2_ release and no O_2_ production as expected; fermentative H_2_ production was only observed in WT cells as reported previously (8-11). Upon illumination, electron flux through PSI was observed accompanied by a sharp CO_2_ uptake due to the carbon concentrating mechanism (CCM) followed by slower CO_2_ fixation via the CBB cycle as also previously observed (Fig. 6) (1). Net O_2_ evolution due to H_2_O splitting at PSII was not detectable immediately upon the start of illumination due to the O_2_ scavenging system of glucose and GOX/Cat. Increasing glucose concentrations successfully delayed the occurrence of net O_2_ evolution (Fig. 7D) and supplementation with the highest concentration of 2 mM glucose kept the cultures anoxic throughout the entire measurements (Fig. 6E, J). Increase of electron flux through PSI parallels the increase of CO_2_ fixation and O_2_ production,when the glucose concentration was insufficient to scavenge all of it (0.85 - 1.5 mM glucose). After 10-15 minutes of light, CO2 fixation ceased due to CO2 limitation in the sample, which was, with a delay of some minutes, accompanied by ceasing O2 production rates (Fig 6A-D, F-I). The electron flow through PSI decreased under CO_2_ limitation and returned to levels comparable to the initial values immediately after onset of illumination.

**Figure 6:**
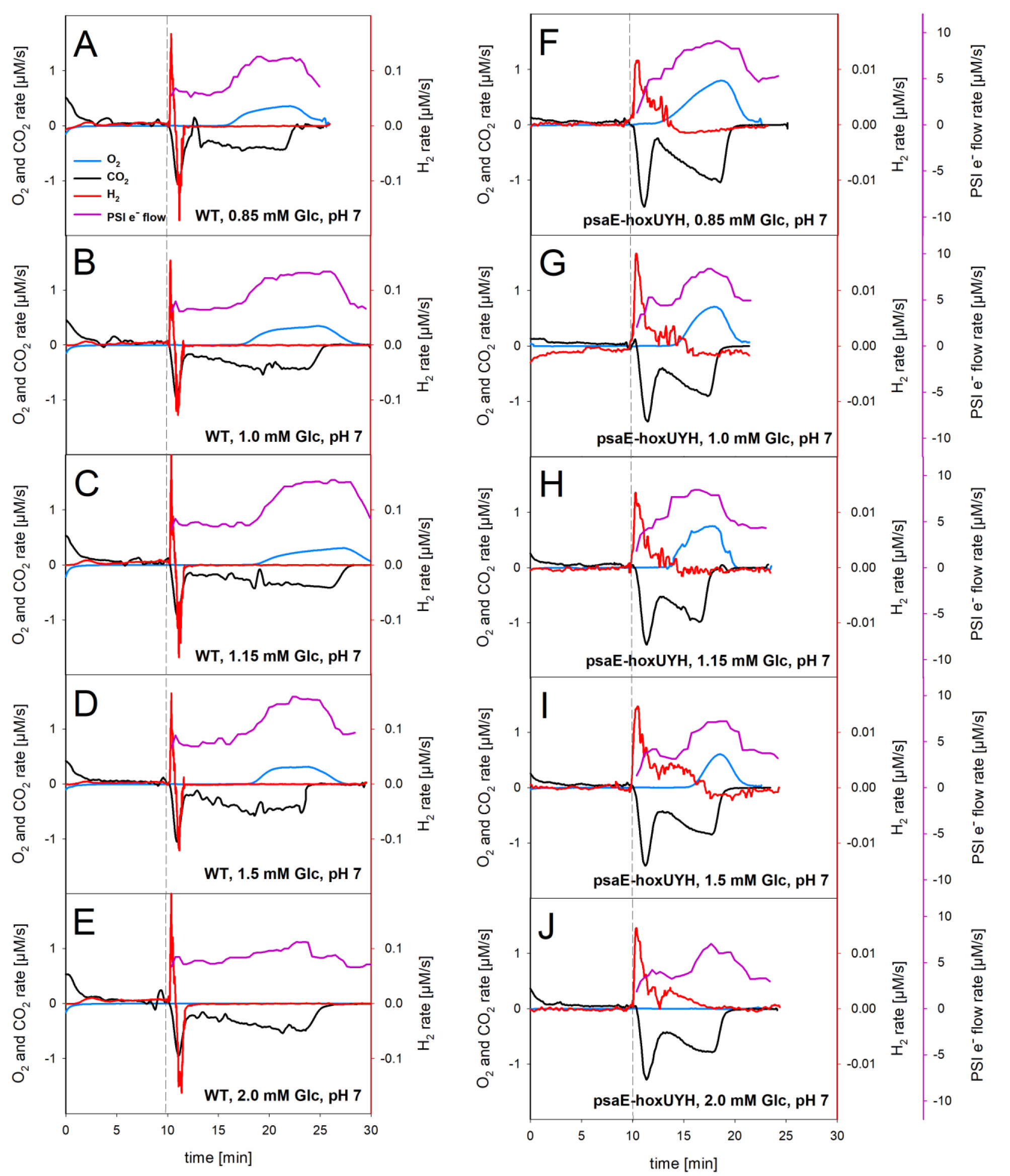
Combined PSI electron flow and simultaneous O_2_, CO_2_ and H_2_ measurements using dark-interval relaxation kinetics (DIRK), membrane-inlet mass spectrometry (MIMS) and Clark-type H_2_-sensor (Unisense). **(A-E)** WT, **(F-J)** psaE-hoxUYH. Cells were adjusted to 20 μg/ml chlorophyll content. Anoxic conditions were achieved with oxygen-scavenging enzyme mix (0.85-2.0 mM glucose, 40 U/ml GOX, 50 U/ml Cat). After 10 min dark adaptation, constant saturating illumination (∼800 μmol photons m^-2^ s^-1^, 625 nm) followed, as indicated with dotted line.

**Figure 7:**
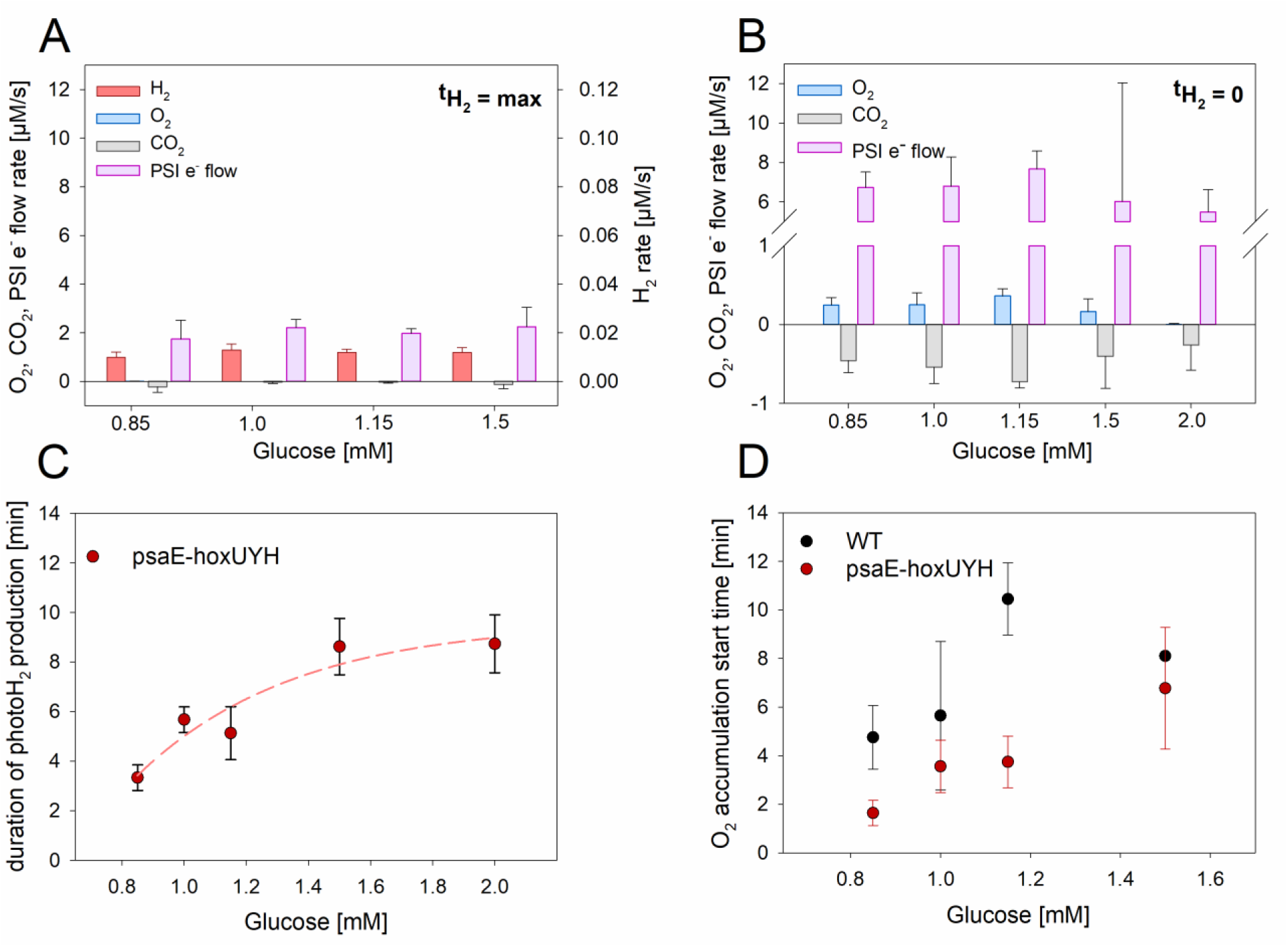
Analysis of combined MIMS, H_2_ and DIRK experiments. **(A)** Rates of O_2_, CO_2_ and PSI e^-^ flux at time of maximal photoH_2_ production rate in psaE-hoxUYH. **(B)** Rates of O_2_, CO_2_ and PSI e^-^ flux at time of termination of photoH_2_ production in psaE-hoxUYH. **(C)** Period of photoH_2_ production in psaE-hoxUYH in relation to added glucose concentration. **(D)** Time of initial O_2_ detection in relation to added glucose concentration in WT and psaE-hoxUYH.

The WT showed the typical short burst of photoH_2_ which was followed by H_2_ consumption. H_2_ uptake occurred almost simultaneously with the CCM. Electron flux via PSI increased during photoH_2_ production and CCM. A delay in O_2_ accumulation due to rising glucose concentrations had no effect on the period of photoH_2_ production in the WT (Fig. 7). In the last minutes of the experiment, CO_2_ fixation ceased and electron flux through PSI was still high (Fig. 6E).

Despite anoxic conditions, the absence of an active CBB cycle as a competitor for electrons and high electron flux through PSI, the WT did not produce photoH_2_ under these conditions. This suggests that another factor, such as the availability of NADH or other yet unknown factors, might limit photoH_2_ production in the WT under these conditions. In psaE-hoxUYH, the initial photoH_2_ rate was about 10 times lower compared to the WT and was not followed by H_2_ uptake. This is expected since H_2_ uptake requires transfer of electrons to Fdx_ox_ and NAD^+^, which in turn requires the HoxE and HoxF subunits, that are absent in the mutant; and a possible electron transfer to the NDH-1 complex would require an interaction between H_2_ase and the complex, which is sterically hindered due to the fusion to PSI. Similar to the WT, electron flux through PSI increased during photoH_2_ production and CCM and further at the onset of CO_2_ fixation via the CBB cycle. The photoH_2_ production rate decreased over a period of several minutes. The accumulation of O_2_ at low glucose concentrations immediately stopped photoH_2_ production in psaE-HoxUYH (Fig. 6F-J). Accordingly, the period of photoH_2_ production could be increased with glucose concentrations up to 1.5 mM. However, a concentration of 2 mM, which kept the cultures anoxic throughout the whole experiment, did not further prolong photoH_2_ production (Fig. 7D). In the absence of O_2_, psaE-hoxUYH was able to produce photoH_2_ in parallel to CO_2_ fixation via the CBB cycle, albeit at lower rates than in the initial phase of illumination. Interestingly, as with WT, experiments using the highest glucose concentrations (2 mM) showed no photoH_2_ production during the final minutes of the experiments, even though net O_2_ production was below the detection limit, CO_2_ fixation had come to a halt, and the electron flow through PSI remained high (Fig. 6 J). Termination of CO_2_ fixation via the CBB cycle did not enhance photoH_2_ production in psaE-HoxUYH (Fig. 6F-J). A comparison of electron flux through PSI at the time of maximum and minimum photoH_2_ production in psaE-hoxUYH shows that PSI electron flux is significantly lower at the timepoint of highest photoH_2_ production (Fig. 7A-B). This indicates that photoH_2_ production in psaE-hoxUYH might not be primarily limited by electron flux through PSI.

### Influence of CO_2_ fixation via the CBB cycle on photoH_2_ production in *Synechocystis* WT and psaE-hoxUYH

To further investigate the influence of CO_2_ fixation as electron competitor for photoH_2_ production in WT and psaE-hoxUYH, a protocol that had previously been established for the green algae *Chlamydomonas reinhardtii* was used. Anaerobic cultures of *C. reinhardtii* were treated with a sequence of 1 s light pulses that alternated with 9 s dark phases to prevent induction of the CBB cycle, thereby eliminating the competition for electrons and allowing continuous photoH_2_ production for a period of up to 70 h (4). In *C. reinhardtii*, photoH_2_ was efficiently produced for 6h, before the rate gradually declined; no CO_2_ was fixed and no biomass was produced during photoH_2_ production (4). *Synechocystis* WT and psaE-hoxUYH were exposed to the described light program consisting of 1 s light and 9 s darkness and supplemented with glucose and GOX/Cat for oxygen consumption. PhotoH_2_ production in the WT could be prolonged to several minutes in comparison to continuous light but eventually switched to H_2_ consumption (Fig. 8). PsaE-hoxUYH, on the other hand, was able to maintain photoH_2_ production for several hours under this light regime, with rates remaining high during the first hours and gradually decreasing over time, as had been observed in green algae (Fig. 8B) (4). The addition of DCMU, which blocks electron transfer between PSII and the plastoquinone pool, reduced photoH_2_ production in WT and significantly lowered photoH_2_ rate in psaE-hoxUYH (Fig. 8A). These observations resemble the photoH_2_ production patterns of WT and psaE-hoxUYH in continuous light, in that photoH_2_ production in WT switches to H_2_ uptake and does not return to photoH_2_ production, whereas photoH2 production in psaE-hoxUYH persists for many minutes or hours, albeit at decreasing rates. CO2 fixation is thus evidently an important, but not the only, limiting factor for photoH2 production in WT and psaE-hoxUYH.

**Figure 8:**
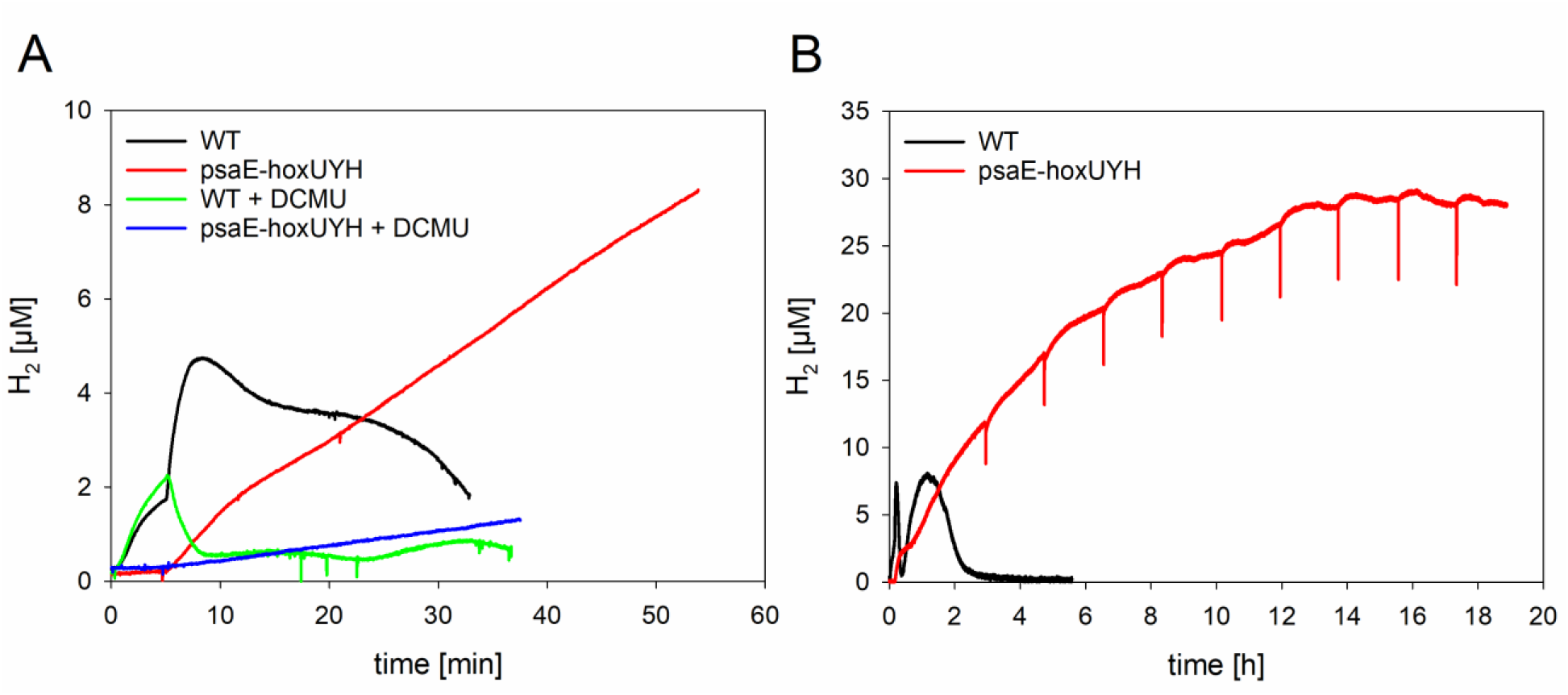
PhotoH_2_ production under a light regime of 1 s light alternating with 9 s darkness in *Synechocystis* WT and psaE-hoxUYH under anoxic conditions (10 mM glucose, 40 U/ml GOX, 50 U/ml Cat) in BG-11_0_, pH 8.0. Cells were adjusted to 20 μg/ml chlorophyll concentration and adapted to darkness for 10 minutes, followed by a light regime of 1 s light (∼800 μmol photons m^-2^ s^-1^, 625 nm) and 9 s darkness over several hours. **(A)** WT and psaE-hoxUYH without and with addition of 40 μM DCMU. **(B)** Long-time measurement of WT and psaE-hoxUYH over 5-18 hours. To prevent sedimentation of cells, a magnetic stirrer was turned on for short time periods several times, leading to appearance of downward signal spikes.

## Discussion

Previously, high yields of 500 μM H_2_ had been reported in psaD-hoxYH cultures under anoxic conditions in continuous light following a lag phase of several hours, although the physiological basis for this H_2_ production was not understood at the time (8). In this study, we have shown that a contaminating bacterium, *Clostridium intestinale*, which possesses several [FeFe]-H_2_ases, is likely responsible for these high H_2_ yields. The O_2_ scavenging system consisting of glucose and GOX/Cat produces gluconate, which cannot be metabolized by *Synechocystis* but serves as a typical carbohydrate source for *C. intestinale (21, 22). Clostridia* are ubiquitous in the environment and the intestines of animals and humans and can survive for years as resistant, dormant spores (23). Gluconate is typically available in the intestines of mammals, which is why many gut bacteria rely on it in combination with the ED pathway (24-26). *C. intestinale*, which was identified in the *Synechocystis* cultures that were kept in a water bath at 28°C (Fig. S3), was previously shown to prefer temperatures between 22 °C to 45 °C and to release CO_2_ and large amounts of H_2_ under fermentative conditions (27, 28). *C. intestinale* as a source for high yields of H_2_ in *Synechocystis* cultures also explains the inconsistent effect of DCMU on H_2_ concentrations that was previously observed (8).

We furthermore show that H_2_ sensors from Unisense (Aarhus, Denmark) react sensitive to light at wavelengths between 400 – 480 nm. For photoH_2_ measurements, polychromatic light should therefore be avoided, whereas monochromatic light at e.g. 625 nm, as used in this study, is suitable. Alternatively, sensor tips can be painted black.

PhotoH_2_ production in the WT and PSI-H_2_ase fusion mutants differs fundamentally. HoxEFUYH requires electrons from NADH from carbohydrate oxidation in combination with Fdx_red_ from PSI. In PSI-H_2_ase fusion mutants, photoH_2_ production should be exclusively based on electrons from photosynthesis. Among the PSI-H_2_ase fusion mutants currently available in *Synechocystis*, psaE-HoxUYH yields the highest amounts of photoH_2_ over the longest production period. Our measurements confirm that CO_2_ fixation via the CBB cycle competes with the H_2_ase for electrons at PSI and thus inhibits or slows down photoH_2_ production even before O_2_-mediated inhibition occurs in WT and psaE-hoxUYH. Similar observations were made for the green algae *Chlamydomonas reinhardtii* (3-5). CO_2_ fixation and presence of O_2_ are limiting factors for photoH_2_ production in both strains. Varying concentrations of glucose were utilized in combination with GOX/Cat to successfully delay the accumulation of O_2_. At the highest glucose concentration (2 mM), no net O_2_ evolution was detected via the MIMS throughout the experiment. PhotoH_2_ production did not continue or resume under anoxic conditions in either the WT or the psaE-hoxUYH strain in the absence of CO_2_ fixation and in the presence of high electron fluxes from PSI. This indicates that additional, yet unknown factors limit photoH_2_ production in WT and psaE-HoxUYH beside CO_2_ fixation and inhibition by oxygen. There might be an unidentified process that might utilize reduction equivalents from cyclic photosynthetic electron transport under these conditions. At the start of illumination, the CBB-cycle is still inactive and CO_2_ concentrations inside the cells are low. Under these conditions, electron acceptors like ferredoxin are instantly reduced, causing an acceptor site limitation at PSI. A transient electron jam occurs, which is mirrored by its low electron transfer rates just at the beginning of the light phase (s. Fig. 6). We hypothesize that prolonged residence time of these electrons at the acceptor side of PSI increases the probability for a transfer to the fused hydrogenase, resulting in the highest photoH_2_ production rates. Subsequent activation of the CBB-cycle and the CCM release this jam and decrease the probability of this transfer and therefore reduce photoH_2_ production to a lower level. Eventually, this low photoH_2_ production ceases completely. Currently, we do not have a conclusive explanation for the halt in photoH_2_ production. One possible explanation might be a build-up of O_2_ below MIMS detection levels that gradually inactivates the H_2_ase. However, the maximum O_2_ concentration inside *Synechocystis* cells that are surrounded by anoxic medium was estimated to be around 0.064 μM (29). H_2_ production by HoxEFUYH was shown to continue at 25-50 % of the maximal rate in the presence of 1 % (10-14 μM) O_2_ and to be reactivated within 90 s by removal of O_2_ and reducing conditions *in vitro* (30). This rather contradicts the idea that small, undetectable amounts of O_2_ could be responsible for halt in H_2_ production. Further experiments are required to clarify this task. Other processes apart from the CBB cycle, such as cyclic electron flow might also compete for electrons with the H_2_ase. An additional fact that has to be taken into account is that not all PSI in our psaE-HoxUYH mutants have HoxUYH fused to them. Electron transfer measurements through PSI do obviously not discriminate between PSI and PSI-HoxUYH which means that the measured fluxes might not reflect the situation that HoxUYH encounters at PSI. In addition, yet unidentified limiting factors might exist. Both in continuous light and in a light regime of 1 s light/9 s darkness, photoH_2_ production turned into H_2_ uptake in the WT. One possible limiting factor might be decreasing NADH and increasing NAD^+^ levels which would shift HoxEFUYH towards H_2_ uptake at high H_2_ concentrations and consequently keep the enzyme unable to produce H_2_, even at high Fdx_red_ levels. PhotoH_2_ production patterns in psaE-hoxUYH differ. In a light regime of 1 s light/9 s darkness, photoH_2_ production occurs at high rates for several hours and thereafter gradually decreases as has been reported for *C. reinhardtii* (4). As the [FeFe]-H_2_ase of *C. reinhardtii* needs only Fdx_red_ as a substrate, the physiological basis of photoH_2_ production in algae and psaE-hoxYH is similar in the sense that both should exclusively rely on photosynthesis. However, both green algae and psaE-hoxUYH show gradually decreasing photoH_2_ production rates over time.

Further studies are required to more precisely determine the limitations of photoH_2_ production in *Synechocystis* WT cells as well as the PSI-H_2_ase fusion mutants.

## Acknowledgements

We thank Viktor Mai and Robert Ehlert for experimental support, Marius Theune for the design of adaptors for photoH_2_ measurements, Anja Heiderich, Heidemarie G_ä_rtner and Alisa L_ä_mmle for technical support and employees from Unisense (Aarhus, Denmark) for technical advice concerning the light sensitivity of H_2_ sensors.

This paper was typeset with the bioRxiv word template by @Chrelli: www.github.com/chrelli/bioRxiv-word-template.

## Author contributions

N.S., M.B., J.A., K.G. conceived and designed the study N.S., F.P., J.R., M.B.,

J.A. performed the experiments, all authors analyzed data, N.S. wrote the first draft, F.P., J.A., K.G. edited, all authors reviewed the manuscript, K.G. acquired funding.

## Materials and Methods

### Construction of Plasmid and Mutant Strain Generation

Gene fragments were amplified with Phusion DNA-polymerase, the respective primers are listed in table S1. Fragments included upstream and downstream recombination sites for the *psaE* locus, *hoxU, psaE, hoxY* and *hoxH* as well as an erythromycin resistance cassette. The psaE-hoxUYH construct was assembled via TAR cloning (12) in pMQ80 (13), which was digested with NaeI and EcoRI beforehand. The protocol was performed as described in Appel et al., 2020. Subsequently, a deletion strain without *psaE* or any *hox* genes (Δ*hox*Δ*psaE*) (10) was transformed with the constructed plasmid. The fusion construct was introduced into the *psaE* locus by homologous recombination.

### Cultivation of *Synechocystis* Strains

*Synechocystis* wild-type and PSI-H_2_ase fusion mutant cells were cultivated under photoautotrophic conditions in BG-11 media (5 mM TES, pH 8.0) at 28°C and 50 μmol photons m^-2^ s^-1^ for three days in 50 ml liquid precultures on an orbital shaker (100 rpm), followed by three days in 200 ml air-bubbled liquid cultures. BG-11 plates and precultures of psaE-hoxYH and psaE-hox-UYH were supplemented with 50 μg ml^-1^ kanamycin and 25 μg ml^-1^ erythromycin. For growth curves, 200 ml cultures were first normalized to an OD_750_ of 0.075.

### Protein Extract Preparation, Gel Electrophoresis and Immunoblotting

Cells were collected by centrifuging and resuspended in ACA buffer (750 mM ε-aminocaproic acid, 50 mM Bis-Tris/HCl, pH 7.0, 0.5 mM EDTA), then broken by vortexing with glass beads (0.17-0.18 μm diameter) for 2 min at full speed at 4°C. Glass beads and cell debris were pelleted by centrifuging at 3000 xg for 10 min. Whole cell extract (WCE) was centrifuged again at 20,000 xg for 20 min to separate crude membranes from soluble protein. The supernatant was kept as the soluble extract, and the crude membrane fraction (CMF) was resuspended in ACA buffer. Chlorophyll concentrations were determined for the WCE and the CMF samples. In the 1D SDS-PAGE immunoblot analysis, an amount corresponding to 10 μg total protein was loaded per lane. These samples were solubilized at room temperature for 1 h in 1x Laemmli sample buffer and separated on 12.5% (w/v) polyacrylamide Bis-Tris gels, using MES running buffer (250 mM MES, 250 mM Tris, 5 mM EDTA, 0.5% (w/v) SDS). For one 1D BN-PAGE gel lane, a CMF sample containing 2 μg chlorophyll was resuspended in 15 μl ACA buffer, and membranes were solubilized for 10 min at 4°C by the addition of 1.66 μl 10% (w/v) β-dodecyl-D-maltoside. Insoluble material was sedimented by centrifugation at maximum speed for 10 min at 4°C in a microfuge. Subsequently, 1.6 μl Coomassie loading solution (750 mM ε-aminocaproic acid, 5% (w/v) Coomassie-G) was added and 18.2 μl was loaded per lane on 5%–12% (w/v) polyacrylamide BN-PAGE 1D gradient gel. For the 2D gels, the protein complexes separated in the first-dimension gel strips were denatured for 1 h in solubilization buffer (66 mM Na_2_CO_3_, 2% (w/v) SDS, 2% (v/v) β-mercapto ethanol, 4 M urea) prior to being layered on 12.5% (w/v) polyacrylamide 6 M urea 2D SDS polyacrylamide gels. All resultant gels were either Coomassie-stained or electroblotted onto a nitrocellulose membrane. For immunoblotting, 5% (w/v) milk powder in 1x PBS-T was used as a blocking solution, and all washing steps were performed with 1x PBS-T. The immunoblot analyses were performed using specific primary antibodies and horseradish peroxidase-conjugated secondary antibodies (anti-rabbit, GE Healthcare). Polyclonal, affinity-purified primary HoxH and HoxY antibodies from rabbit raised against full-length *E. coli* over-expressed protein were used (14). The PsaB, PsaC and PsaD antibodies were purchased from Agrisera (Umeå) (15).

### Hydrogen Measurements

Cells were grown under photoautotrophic conditions as described above, sedimented (5 min, 8000 xg, RT) and resuspended in BG-11_0_ (BG-11 without NaNO_3_) with either pH 7.0 or 8.0. Hydrogenase activity assays were performed with a Clark-type disc electrode S1 (Hansatech Instruments) as described previously (16), using 10 mM sodium dithionite and 5 mM methyl viologen.

Fermentative H_2_ and transient photoH_2_ production were monitored with Clark-type H_2_ sensors (Unisense, Aarhus, Denmark). The chlorophyll concentration of samples was adjusted to 20 μg/ml. For O_2_ scavenging, GOX/Cat (40 U/ml glucose oxidase, 50 U/ml catalase) and 0.85-10 mM glucose were added. Cells were measured for 10 minutes in darkness, followed by illumination with monochromatic light at 625 nm and ∼800 μmol photons m^-2^ s^-1^ (MULTI-COLOR-PAM, Walz, Effeltrich, Germany) to induce photosynthesis.

Long-time H_2_ experiments were carried out in a MicroRespiration System (Unisense, Aarhus, Denmark) consisting of a rack with magnetic stirrers, double glass chambers and O_2_/H_2_-sensors which were set up in a water bath for temperature control (28°C) (Fig. S3). Each chamber was filled with 13 ml cell suspension normalized to a chlorophyll concentration of 20 μg/ml and supplied with GOX/Cat (40 U/ml glucose oxidase, 50 U/ml catalase) for anoxic conditions. Cells were adapted to darkness for one hour, followed by illumination with ∼800 μmol photons m^-2^ s ^-1^ polychromatic light (12P HEX, ADJ, Thomann, Burgebrach, Germany).

### Membrane Inlet Mass Spectroscopy (MIMS)

Simultaneous O_2_ and CO_2_ measurements were performed using the DELTA™ Q Isotope Ratio Mass Spectrometer (Thermo Scientific™). Cells were prepared as described in hydrogenase activity assays and resuspended in BG-11_0_ (50 mM MOPS, pH 7.0). The lower pH buffer was chosen to reduce the noise of the CO_2_ signal that occurs at pH 8.0. Additionally, the lower pH shifts the equilibrium of CO_2_ and HCO_3-_, decreasing the concentration of dissolved inorganic carbon to a level that is consumed within 30 minutes. Carbonic anhydrase (400 U/ml) was added to ensure the equilibrium between CO_2_ and HCO_3-_. The argon trace recorded simultaneously was used to reduce the noise and to correct for the gas consumption by the instrument.

### Dark Interval Relaxation Kinetics (DIRK)

For investigations of electron flux through PSI, cells were prepared according to the protocol of transient photoH_2_ measurements. Dark interval relaxation kinetics were measured using a near-infrared spectrometer DUAL-KLAS/NIR (Walz) as described in Theune et al. (2021).

### Metagenome Analysis

Samples from lasting H_2_ measurements were analyzed as raw paired-end Illumina reads (2x 150 bp) obtained from Eurofins, generated using extracted genomic DNA and sequenced on an Illumina NovaSeq 6000.

Bioinformatical analysis of the Illumina sequencing data was carried out using a workflow in the open-source European Galaxy server (https://usegalaxy.eu; accessed in October-November 2024). The workflow ran quality control and initial sequence filtering. Paired end FASTQ reads were combined using FASTQ Interlacer (Galaxy Version 1.2.0.1+galaxy0). Taxonomic profiling analysis was carried out using MetaPhlAn2 set to search taxons bacteria and eukaryotes with varying statq values to estimate the noise.

To amplify *Clostridium intestinale* [FeFe]-hydrogenase genes directly from contaminated culture samples, Phire Plant Direct PCR Master Mix (Thermo Scientific™) was used according to its manual. For this, 1 μl cells were added in 20 μl Master Mix and initial denaturation was prolonged to 5 minutes for sufficient cell lysis.

