## Supplementary Material for "Physiological basis of photosynthetic hydrogen production in the cyanobacterium *Synechocystis*"

### Content

**Table S1.** List of primers for construction of psaE-hoxUYH plasmid

**Figure S1.** Light sensitivity of Unisense $H_2$-sensors in long-time photo$H_2$ experiments.

**Figure S2.** Sample collection for metagenome analysis during long-time $H_2$ and $O_2$ measurements of Δ*hox* under anoxic conditions.

**Figure S3.** Experimental set up of long-time $H_2$ measurements.

**Figure S4.** Replicate (n=4-5) representation of combined PSI electron flow and simultaneous $O_2$, $CO_2$ and $H_2$ measurements using dark-interval relaxation kinetics (DIRK), membrane-inlet mass spectrometry (MIMS) and Clark-type $H_2$-sensor (Unisense).

**Table S1.** List of primers for construction of psaE-hoxUYH plasmid

| Primer name | sequence | Fragment amplified |
|---|---|---|
| Eco-PsaEout1 | ACTCTCTACTGTTTCTCCATACCCGTTTTTTTGG<br>GCTAGCTACCTCATGTCTCTTTGCTCACCAT | 5´ part of *psaE*<br>(upstream recombination site) |
| HoxUN-PsaEin1 | AATGGTTAAAGTAACAACAGACATGATTTAATTC<br>CTTAGATAATTGCGA | |
| PsaEin1-HoxUN | TCGCAATTATCTAAGGAATTAAATCATGTCTGTT<br>GTTACTTTAACCATT | *hoxU* |
| PsaEN-HoxUC16 | TTTTCAGTACTTTTACTTTCACTACCACTACCAC<br>TACTTTTTTCCCTGACCCATTCCTTTTCTTTA | |
| HoxYC-PsaEN16 | AAAGTAGTGGTAGTGGTAGTGAAAGTAAAAGTAC<br>TGAAAAAAGTGGTGACAAAGTTAGAATCAAACGC | *psaE* |
| HoxYN-PsaEC | AACGAATTTTAGCCATGATTAAAATCTCCTAGAC<br>TATTTTGCCGCCGCTTGCACCAAT | |
| PsaEC-HoxYN | ATTGGTGCAAGCGGCGGCAAAATAGTCTAGGAGA<br>TTTTAATCATGGCTAAAATTCGTT | *hoxY*+*hoxH* |
| Em-HoxH | TAATTTCTTTTTTCGTCGACCGTCTGAATGTTTT<br>TTGTTTAATCCC | |
| HoxH-Em | AAACAAAAAACATTCAGACGGTCGACGAAAAAAG<br>AAATTAGATAAA | Erythromycin-cassette |
| PsaEin2-Em | TGCAAATTTAGTCCACAGTCAAGAGTCGACTTAC<br>TTATTAAATAATT | |
| Em-PsaEin2 | AATTATTTAATAAGTAAGTCGACTCTTGACTGTG<br>GACTAAATTTGCA | 3´ part of *psaE*<br>(downstream recombination site) |
| NaeI-PsaEout2 | CGAAGCAGGGTTATGCAGCGGAAAGTATACCTTA<br>ACCGCCAGCCTGCCACCAACCGTTGATCCA | |

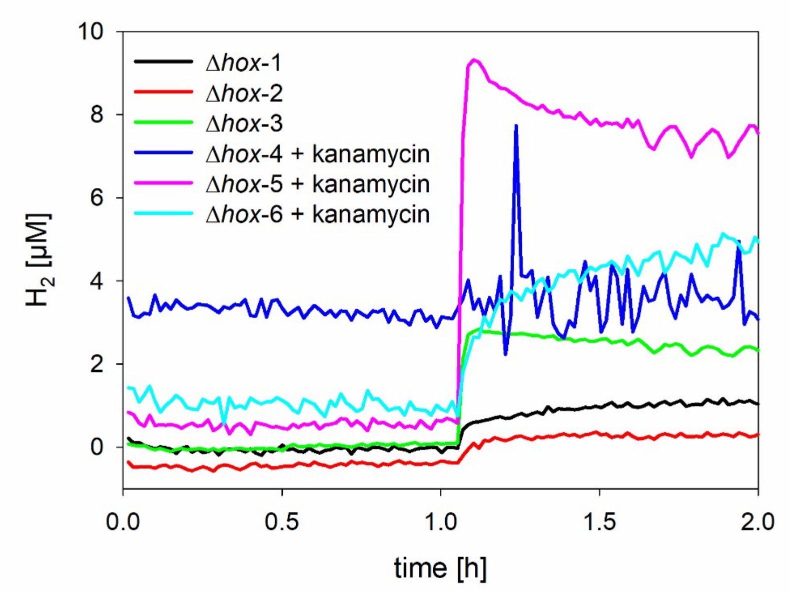


**Figure S1.** Light sensitivity of Unisense $H_2$-sensors in long-time photo$H_2$ experiments from Fig.2B. Cells were adapted to darkness for one hour. The arrow indicates when light (~800 µmol photons $m^{-2}$ $s^{-1}$, 12P HEX, ADJ) was switched on. Six sensors show different sensitivity corresponding to signals of up to 9 µM $H_2$.

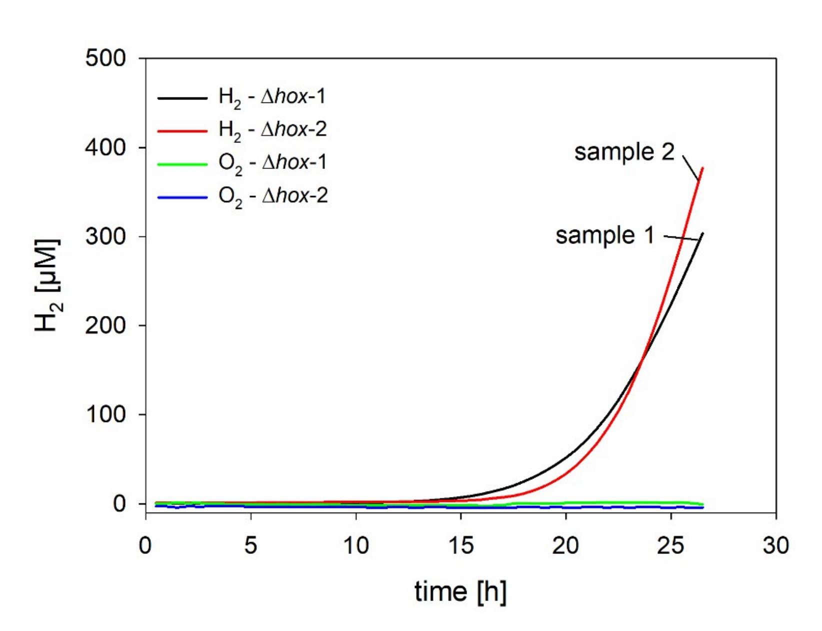


**Figure S2.** Sample collection for metagenome analysis during long-time $H_2$ and $O_2$ measurements of Δ*hox* under anoxic conditions (10 mM glucose, 40 U/ml glucose oxidase, 50 U/ml catalase). Cells were adapted to darkness for one hour, followed by constant illumination (~800 µmol photons $m^{-2}$ $s^{-1}$, 12P HEX, ADJ). Samples were taken during high $H_2$ production rate phases, after which the experiment was stopped.

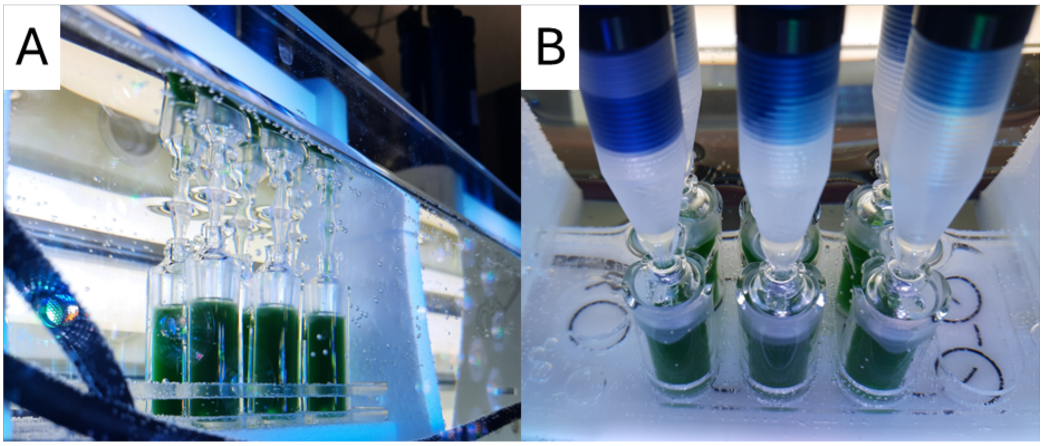


**Figure S3.** Experimental set up of long-time $H_2$ measurements. Cells were transferred to gas tight glass double chambers, placed in a MicroRespiration Rack containing a magnetic stirrer (Unisense, Aarhus, Denmark). Sensors were inserted into the chambers from the top. The rack was placed in a water bath to maintain a temperature of 28°C, thus the sensor tips might introduce some water into the samples during insertion. (A) Side-view, (B) top-view.

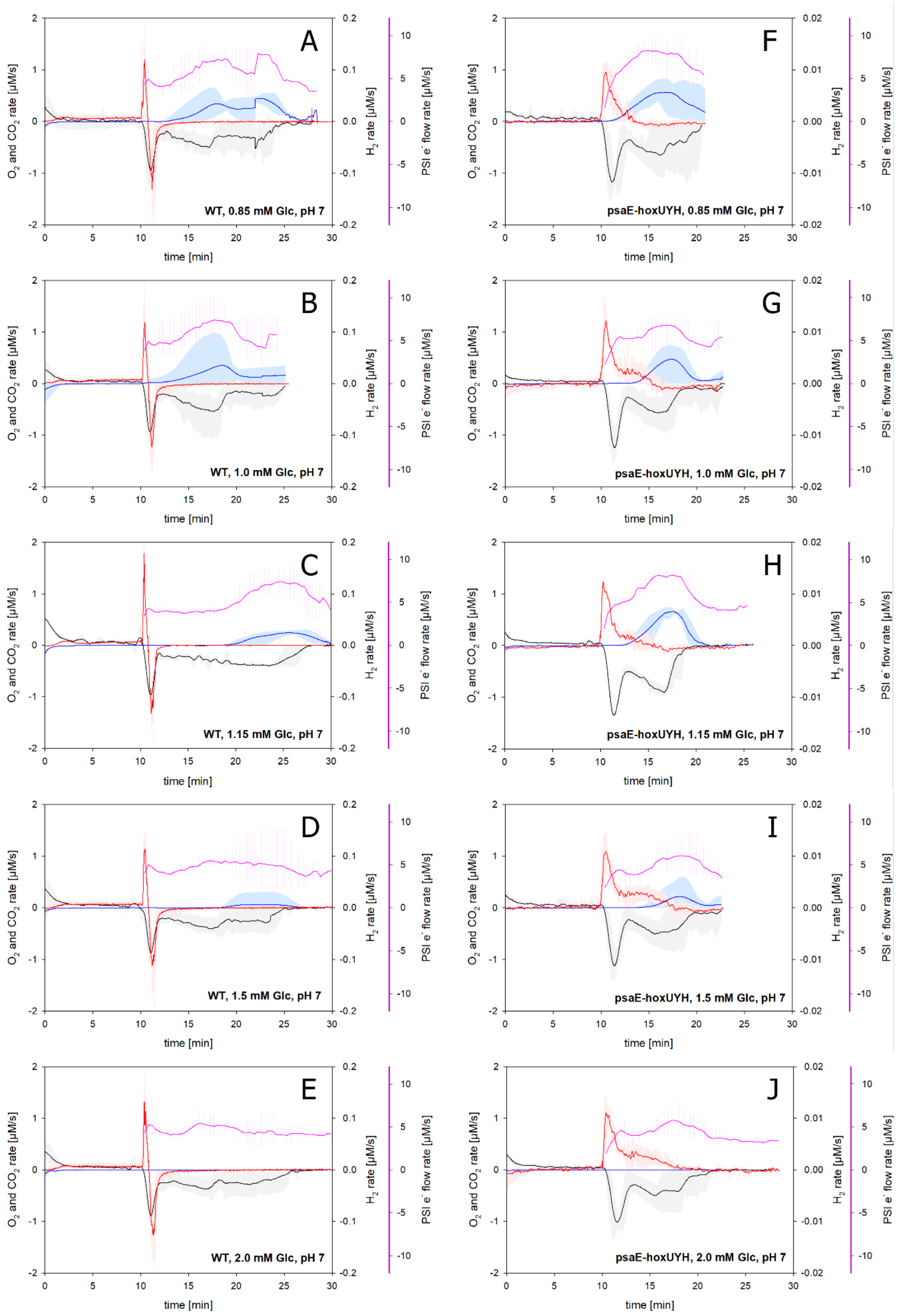


**Figure S4.** Replicate (n=4-5) representation of combined PSI electron flow and simultaneous $O_2$, $CO_2$ and $H_2$ measurements using dark-interval relaxation kinetics (DIRK), membrane-inlet mass spectrometry (MIMS) and Clark-type $H_2$-sensor (Unisense). (A-E) WT, (F-J) psaE-hoxUYH. Cells were adjusted to 20 µg/ml chlorophyll content. Anoxic conditions were achieved with oxygen-scavenging enzyme mix (0.85-2.0 mM glucose, 40 U/ml glucose oxidase, 50 U/ml catalase). After 10 min dark adaptation, constant illumination (~800 µmol photons $m^{-2}$ $s^{-1}$, 625 nm) followed. Due to differing periods of $CO_2$ fixation in experiments, some respective curves show a step-like appearance.
